# Two-Dimensional Hermite Filters Simplify the Description of High-Order Statistics of Natural Images

**DOI:** 10.1101/060947

**Authors:** Qin Hu, Jonathan Victor

## Abstract

Natural image statistics play a crucial role in shaping biological visual systems, understanding their function and design principles, and designing effective computer-vision algorithms. High-order statistics are critical for conveying local features, but they are challenging to study – largely because their number and variety is large. Here, via the use of two-dimensional Hermite (TDH) functions, we identify a covert symmetry in high-order statistics of natural images that simplifies this task. This emerges from the structure of TDH functions, which are an orthogonal set of functions that are organized into a hierarchy of ranks. Specifically, we find that the shape (skewness and kurtosis) of the distribution of filter coefficients depends only on the projection of the function onto a 1-dimensional subspace specific to each rank. The characterization of natural image statistics provided by TDH filter coefficients reflects both their phase and amplitude structure, and we suggest an intuitive interpretation for the special subspace within each rank.

## 1. Introduction

Achieving a thorough understanding the statistics of our visual environment is important from both a biological point of view and an engineering point of view. The biological relevance is that the statistics of the natural environment are a strong constraint under which visual systems evolve,develop, and function[1]. The engineering relevance is that a knowledge of image statistics is important for many problems in computer vision [2], including image de-noising, image classification [3–6]), image compression, and texture synthesis [7]. However, understanding image statistics is hampered by the simple fact that the space of image statistics is so large. Here we describe some progress in this direction: a specific filter-based approach that identifies a hidden symmetry, providing a simplified description of high-order natural image statistics, specifically,those of order three and four.

The reason for our focus on high-order statistics is that they carry local visual features, such as lines, corners, and edges [8, 9], but – because of the curse of dimensionality, they are challenging to analyze. In contrast, second-order statistics are concisely captured by the power spectrum, because it is the Fourier transform of the autocorrelation function. As is well-known, the power spectrum of natural images is approximately *k*^-2^ (where *k* is spatial frequency) [10, 11]. However, while the power spectrum captures important spatial regularities of natural images – such as distance-independent scaling [12], it is far from a complete statistical description of natural images. For example, a synthetic image consisting of Gaussian noise with a *k*^−2^ power spectrum looks drastically different from a real natural image, even though the spectra are similar. Conversely modifying a natural image by flattening its power spectrum but preserving its phases leaves its salient spatial features readily recognizable. Thus, most of the features that make an image look “natural,” such as edges and contours, are coded in its phases as well as its Fourier amplitudes [8, 9, 13]. Translated into the spatial domain, these phase correlations correspond to image statistics that are ignored by the power spectrum: joint distributions of image intensities at three or more points, and aspects of the pairwise intensity distributions beyond their variances and covariances.

Since a direct tabulation of the joint distribution of multiple pixel values is impractical, a natural strategy is to focus on specific univariate distributions – namely, the distribution of outputs of filters (“filter coefficients”) placed on images. Typically, this approach is implemented with filter profiles that have a prominent orientation and dominant spatial frequency – either Gabor functions or Gabor-like wavelets, a choice motivated by concepts of visual processing and independent components analysis of natural images[14, 15]. For natural images, the distributions of wavelet coefficients are highly kurtotic, having sharp peaks and much longer tails compared to a Gaussian distribution with the same variance [16]. Interestingly, [3] showed that this could be used to distinguish natural images from synthetic ones (including realistic computer-generated scenes), by applying linear classifiers to a feature space of wavelet coefficients. Other investigators have also used wavelet coefficients as a starting point, but focused on the extent to which wavelet coefficients are independent [17, 18]. Thus, the filter approach provides a useful characterization of natural image statistics -- but even with a filter-based approach, the number of parameters required to describe high-order image statistics is still large. Since a two-dimensional basis set is a two-parameter family, the number of parameters required to specify these filter coefficient distributions is still quite large.

Here we show that the description of these filter coefficient distributions is simplified when, instead of Gabor-like filters, we use a set of filters that have a high degree of symmetry. These filter functions – the two-dimensional Hermite functions (TDH’s) [19–25]-- form an orthonormal basis that is halfway between the pixel basis and the Fourier basis, and their shapes are quite different from that of Gabor-like filters or one-dimensional wavelets. Although the TDH functions form a full two-dimensional basis set, we find that the distribution of filter coefficients for natural scenes depends chiefly only a single parameter, their rank. The nature of this simplification reflects both the phase and amplitude characteristics of natural scenes.

## 2. Materials and Methods

### 2.1. Two-dimensional Hermite functions: definition and properties

We analyze image statistics via the distribution of values that result from filtering them with two-dimensional Hermite (TDH) functions. TDH’s (Figure 1) are a set of two-dimensional functions consisting of a product of Hermite polynomials multiplied by a Gaussian envelope. Like wavelets, they are filter functions that are limited in space and spatial frequency. However, they several other mathematical properties, including additional symmetries. First, the TDH’s are optimally symmetrical with respect to space and spatial frequency: other than a multiplicative constant, each TDH is its own Fourier transform. Second, they are orthonormal functions, and as a set, form a complete basis set for functions of two variables. Third, the TDH’s are grouped into “ranks”: the sole member of the zeroth rank is an ordinary Gaussian; higher rank ranks contain functions of increasing spatial complexity. Finally, within each rank, the TDH’s have an extended steerability property. This includes ordinary steerability – the filters can be rotated by forming simple linear combinations – but also, linear combinations within rank provide equivalent basis sets that are separable in Cartesian coordinates (see rows of Figure 1).

**Figure 1.**
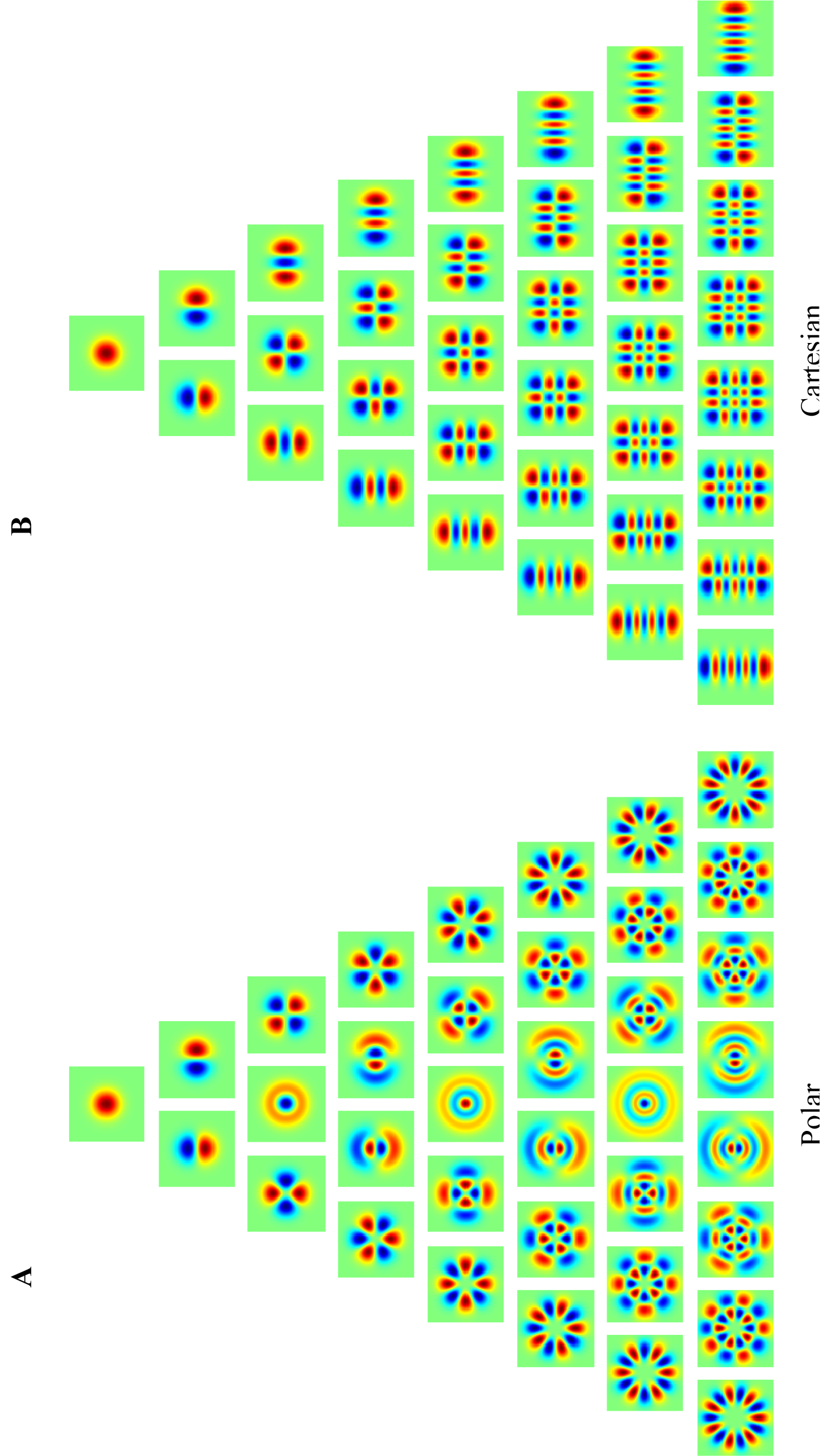
Two-dimensional Hermite (TDH) functions of rank 0 to 7, in (A) polar form and (B) Cartesian form. The pseudocolor scale (red positive, blue negative) is chosen separately for each function to cover the entire range. (Modified from Figure 1 in [12], Victor et al. 2006, J. Neurophysiol., Am Physiol Soc, with permission.)

Below we define these functions in abstract terms and then give their explicit polynomial expansions; the former makes these and other key properties transparent, while the latter is necessary for computation. For further details on this approach, see [25]; other descriptions of the properties of these functions in the context of image processing may be found in [19–24].

Taking inspiration from [26, 27], we define the TDH’s as the eigenvectors of the operator *D*^1/2^*BD*^1/2^, where *D* consists of spatial windowing by a two-dimensional Gaussian function (i.e., pointwise multiplication), and *B* consists of filtering by a two-dimensional Gaussian spatial frequency window (i.e., pointwise multiplication in the spatial frequency domain). This operator *D*^1/2^ *BD*^1/2^ is self-adjoint, and has a discrete set of eigenvalues [25]. The approach of [28] shows that these are of the form *λ* = *η*^1+*r*^, for a positive constant *η* < 1, where the rank, *r*, ranges over the non-negative integers [25]. It also shows that the *r* th rank contains *r* +1 linearly independent functions [25].

Since *D* corresponds to confinement in space, and *B* corresponds to confinement in spatial frequency, a TDH function *f* has the property that successive windowing in space and spatial frequency results in multiplication by a constant (the eigenvalue *λ D*^1/2^ *BD*^1/2^ *f* = *λf*. That is, for functions *f* corresponding to eigenvalues *λ* close to 1, these windowing operations have a small effect – which formalizes the notion that *f* is confined in both space and spatial frequency.

Since the eigenvalues are all of the form *λ* = *η*^1+*r*^, the TDH function of rank *r* = 0 has the eigenvalue that is closest to 1, and is therefore the most confined. Successive ranks have exponentially-declining eigenvalues, and are therefore progressively less confined (i.e., is more extensive spatially and contains a progressively broader range of spatial frequencies). TDH functions at different ranks are orthogonal, since they correspond to different eigenvalues of the self-adjoint operator *D*^1/2^ *BD*^1/2^.

The extended steerability of the TDH functions is a consequence of combining this setup with the fact that a circularly-symmetric Gaussian is separable both in Cartesian and polar coordinates. As a consequence, both *D* and *B* have polar symmetry and separability in Cartesian coordinates, These symmetries are inherited by *D*^1/2^*BD*^1/2^ as well, and must be retained by the eigenspaces, so the existence of Cartesian and polar-symmetric eigenvectors are guaranteed. Since any set of *r* +1 linearly independent eigenvectors forms a basis for each rank, it follows that we can express the Cartesian and polar basis sets as linear combinations of each other.

### 2.2. Two-dimensional Hermite functions: explicit expressions

As described in §2.1, there are two natural basis sets for the TDH functions of rank *r* : polar and Cartesian. The polar basis functions are specified by their rotational symmetry (an integer *μ*, for which a rotation by *2π / μ* leaves the function unchanged) and the number of zero-crossings along each radius (an integer *ν*). These indices are related to the rank *r* by *r = μ + 2V*. For *μ* > 0, the basis functions form “cosine” and “sine” pairs:

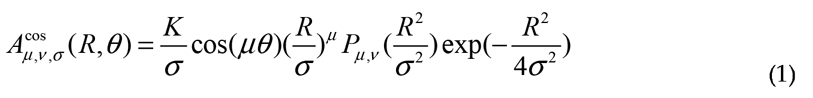

and

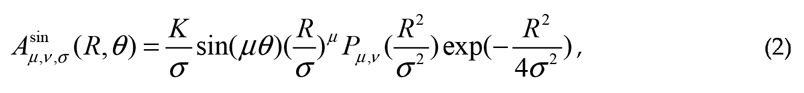

where *σ* sets the overall size of the filter set, *K* is a normalization constant, and *P_μ,v_*(*u*) is a radial polynomial defined by

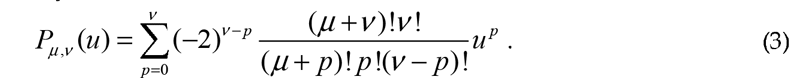

For each even ranks, there is also an unpaired basis function, corresponding to *μ* = 0 and *V = r* / 2. These basis functions have no angular dependence (central column of Figure 1A), and are given by 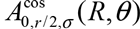

A typical Cartesian basis function has the appearance of vignetted (*j* +1)×(*k* +1) checkerboard, where there are *j* vertical zero-crossings, *k* horizontal crossings, and these indices are related to the rank by *r = j + k*. It is given by

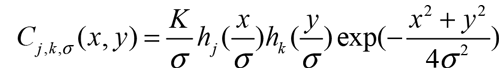

where *h_j_*(*u*) and *h_k_*(*u*) are Hermite polynomials, normalized so that they have the generating function

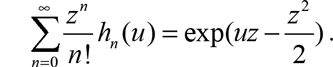

As detailed in §2.4, we calculate image statistics of natural images filtered by the polar TDH’s, and then use steerability to calculate the statistics of images filtered by other TDH’s of a given rank, including the Cartesian TDH filters (as indicated in Figure 2) and intermediate ones. Note that this “steerability” is much more than geometric rotation, as it allows for filters of different shapes and symmetries (see Figure 4 below) to be represented in terms of a small basis set.

**Figure 2.**
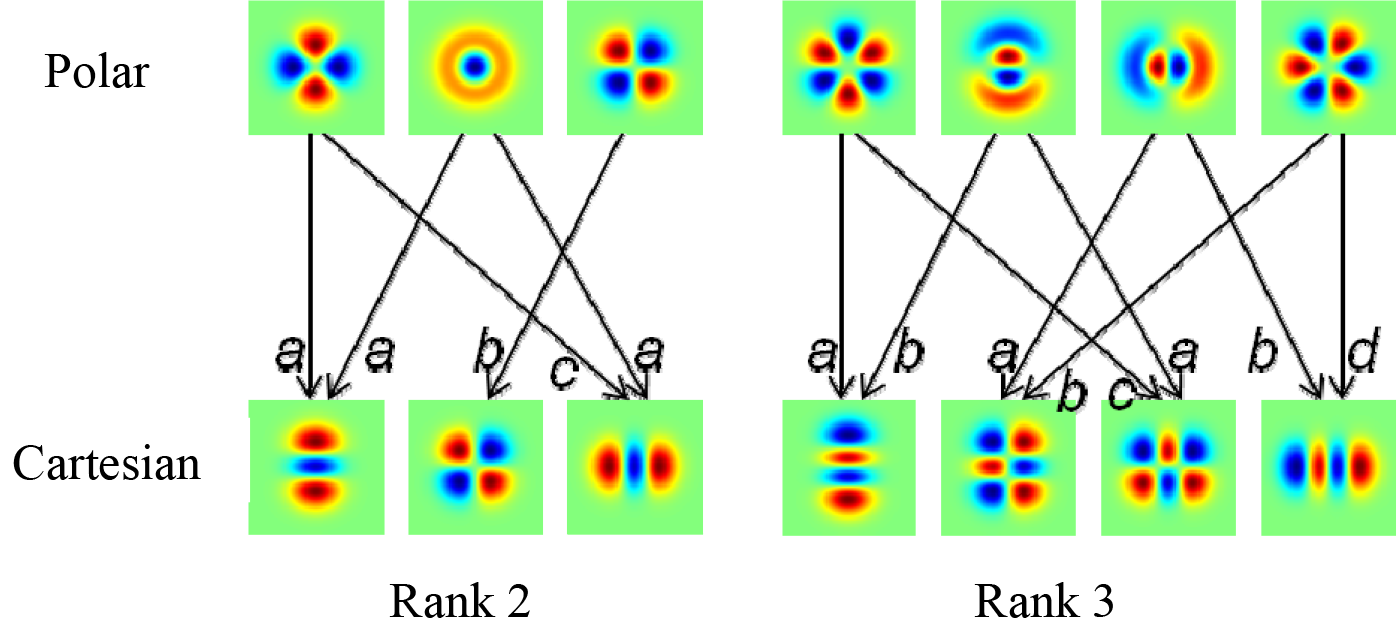
Cartesian TDH functions are linear combination of polar TDH functions. Examples are shown for rank 2 (left) and rank 3 (right). For rank 2, the coefficients are 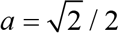, *b* = 1, 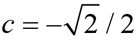. For rank 3, the coefficients are *a* = 1/2,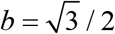, 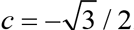, *d* = −1/2.

### 2.3. Natural images

All 4167 images from the van Hateren natural image database [15] (van Hateren & van der Schaaf, 1998) were chosen for analysis. Each image is 1536 by 1024 pixels, with each pixel’s intensity represented by a 16-bit unsigned integer. The images mainly contain landscapes and plants, but occasionally manmade objects such as houses appear.

### 2.4. Analysis

To characterize high-order statistics of natural images, we calculated the skewness and kurtosis (as “excess kurtosis“) of the distribution of filter coefficients, i.e., the distribution of values that result from convolving the images with TDH functions. To focus on the structure of the individual scenes (rather than the overall differences across scenes), skewness and kurtosis were calculated individually for each image, and values were then averaged across the image database.

As shown in Figure 3, this calculation was carried out across 7 spatial scales, spaced in approximately octave steps. The smallest scale used *σ* = 7/12 (0.58) pixels and the largest, *σ* = 511/12 (42.6) pixels. At each scale, the image was convolved with polar TDH functions of ranks 0–7 (36 filters in all), and the convolution was sampled at points placed in a rectangular grid on the filtered image. Filters centers were separated by 10 pixels for scales 1–5 and 50 pixels for scales 6 and 7. We then calculated the pure and mixed moments of these distributions up to order 4, and used the extended steerability property (detailed below) to go from the moments for the polar TDH functions to the moments for arbitrary TDH functions. From these moments, skewness and kurtosis were then calculated in the standard fashion.

**Figure 3.**
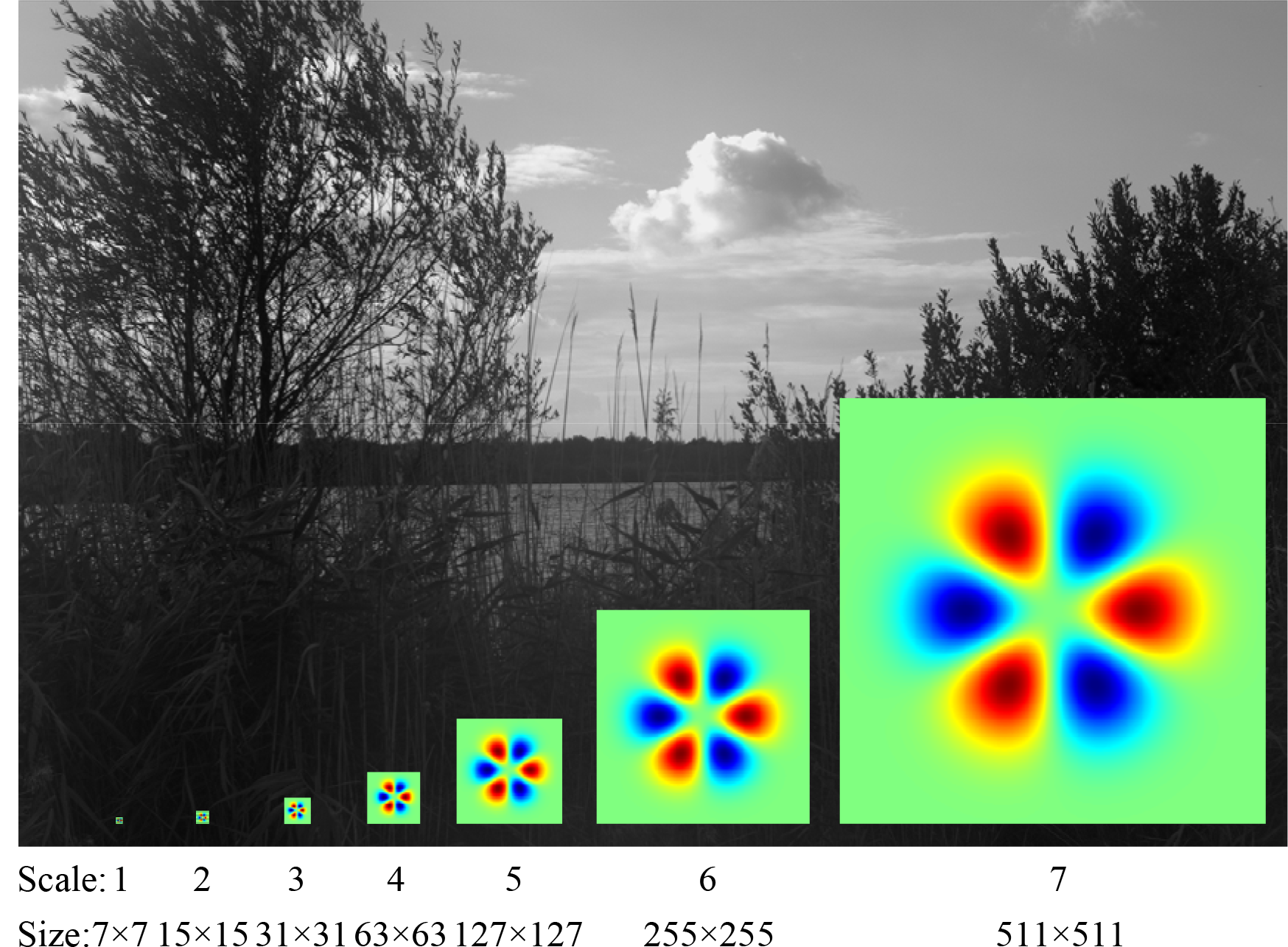
The seven filter sizes used to calculate image statistics, compared to the size of natural images used in this study (1536×1024).

In detail, computation of the skewness and kurtosis for all TDH functions *F* of rank *r* were carried out in parallel, as follows. For each image *I*, we calculated the pure moments for each polar basis function *f*

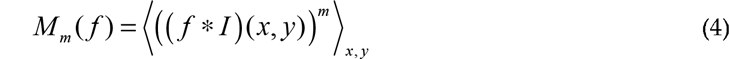

up to *m* = 4, and, analogously, the mixed moments for each pair of functions *f* and *f′*

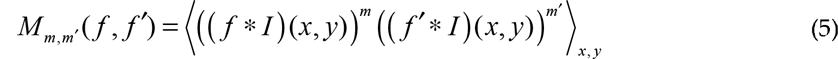

up to *m* + *m′* = 4, and, analogously, the mixed moments *M*_1,1,1_(*f*, *f′*, *f″*), *M*_2,1,1_(*f*, *f′*, *f″*) and *M*_1,1,1,1_(*f*, *f′*, *f″*, *f‴*).

To use the steerability property, we wrote the filter function *F* as a linear combination of the polar basis functions of that rank:

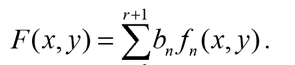

Therefore, the convolution of *F* with an image *I* can be calculated as a linear combination of the convolutions of the basis functions with the image,

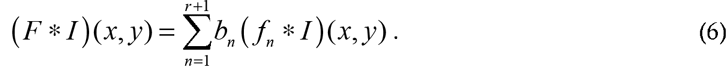

Expressions relating the moments of the distribution of the filter coefficients for *F* to the moments for the basis functions *f_n_* now follow via multinomial expansion of (6), using (4) and (5):

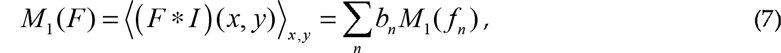

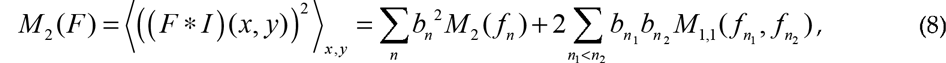

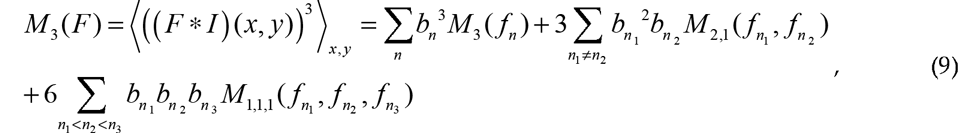

and

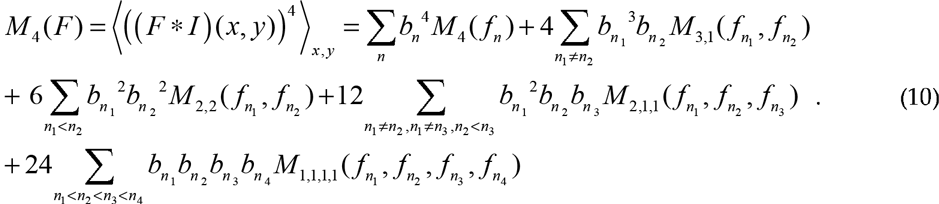

As is standard, the cumulants of the distribution of the filter outputs of *F* are determined from its moments by

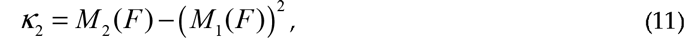

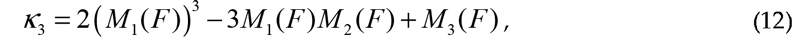

and

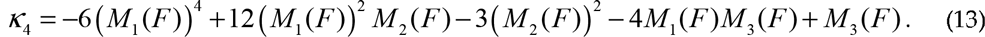

Skewness and (excess) kurtosis are ratios of the cumulants:

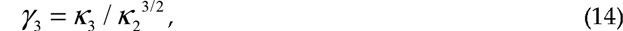

and

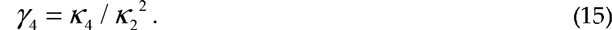

## 3. Results

We characterized the high-order statistics of natural images via the distribution of filter coefficients for two-dimensional Hermite (TDH) functions. We present the findings for rank 2 first because this low rank allows for a detailed visualization, and then turn to higher ranks.

### 3.1. Statistics of rank-2 TDH filter coefficients for natural images

To visualize the results for rank 2, we note that the full set of rank-2 filters can be regarded as points on the surface of an ordinary sphere (Figure 4). This follows from the general observation that the *r* th rank of TDH functions is spanned by contains *r* +1 orthonormal filters, so the full set of unit-magnitude filters of rank *r* (i.e., the full set of unit-magnitude linear combinations of these *r* +1 basis elements) may be regarded as the surface of a sphere in (*r* +1)-space. In this spherical representation of rank-2 TDH functions shown in Figure 4, the polar filters correspond to one set of orthogonal directions, the Cartesian filters to a second orthogonal set of directions, and intermediate directions correspond to mixtures of polar or Cartesian filters. The latitude (altitude) indicates the size of the projection onto the target-like TDH function. For TDH functions at the same latitude, the azimuth on the sphere corresponds to the orientation (i.e., the in-plane rotation angle) of the filter function.

**Figure 4.**
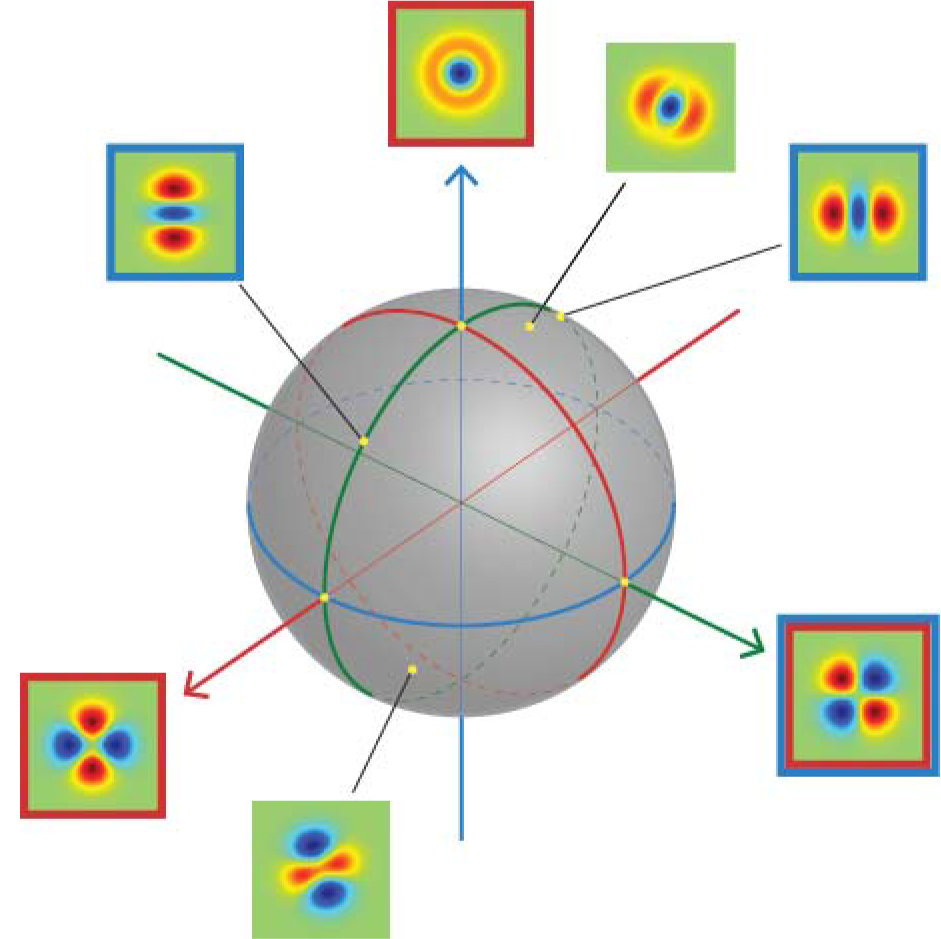
Generalized steerability of the rank 2 TDH filters. Each unit-magnitude filter corresponds to a point on the surface of a sphere. The polar and Cartesian basis functions form two sets of orthogonal coordinate axes Filters with a red frame are polar TDH filters; filters with blue frame are Cartesian TDH filters; one filter is in both sets as indicated by its two frames. Filters without frame are intermediate filters; they can be constructed from a linear combination of either polar or Cartesian filters.

Figure 5 shows skewness and kurtosis of the distributions for all TDH filters of rank 2, plotted on the filter space shown in Figure 4. Skewness and kurtosis depends strongly on latitude, but is largely are largely independent of orientation, although there is a small dependence of kurtosis at orientation at the two largest scales. Skewness is largest for the circularly-symmetric (target-like) filters at the poles and zero for filters on the equator, while kurtosis was smallest for the target-like filters, and largest for filters on the equator.

**Figure 5.**
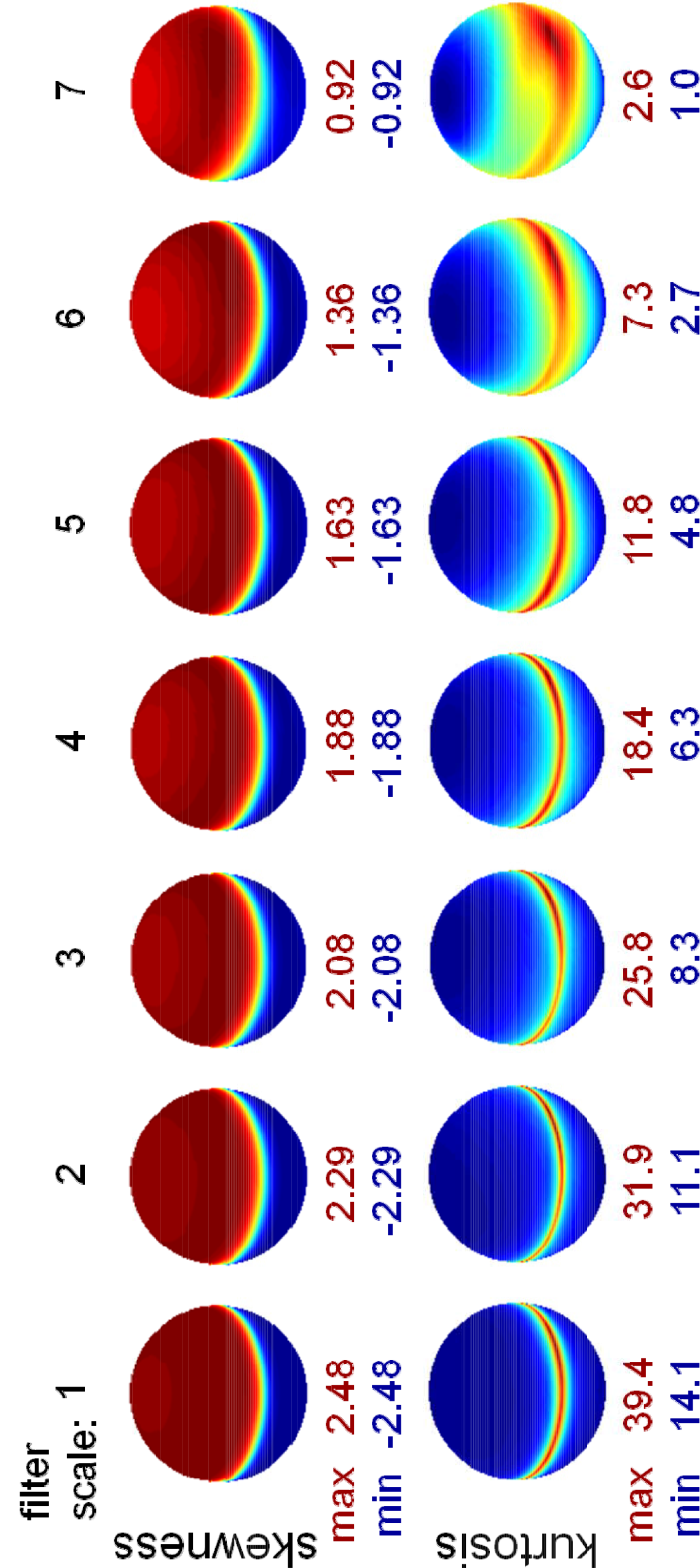
Skewness and kurtosis for natural images filtered by rank 2 TDH filters across 7 spatial scales. Each sphere represents the filter space of unit-length rank 2 TDH filters (oriented as shown in Figure4). Skewness and kurtosis are averaged across all filtered images, and plotted as a function of direction in the filter space. The pseudocolor scales for each skewness and kurtosis map are set to range from blue (minimum) to red (maximum). The minimum and maximum skewness and kurtosis values are shown under each sphere.

### 3.2. Statistics of higher-rank TDH filter coefficients for natural images

For higher ranks, a similar visualization strategy is not possible, so we begin with the skewness and kurtosis for each of the filters in the polar basis set (Figure 6). We focus on filter scale 4, the middle of the range studied; other filter scales gave a similar pattern of results.

**Figure 6.**
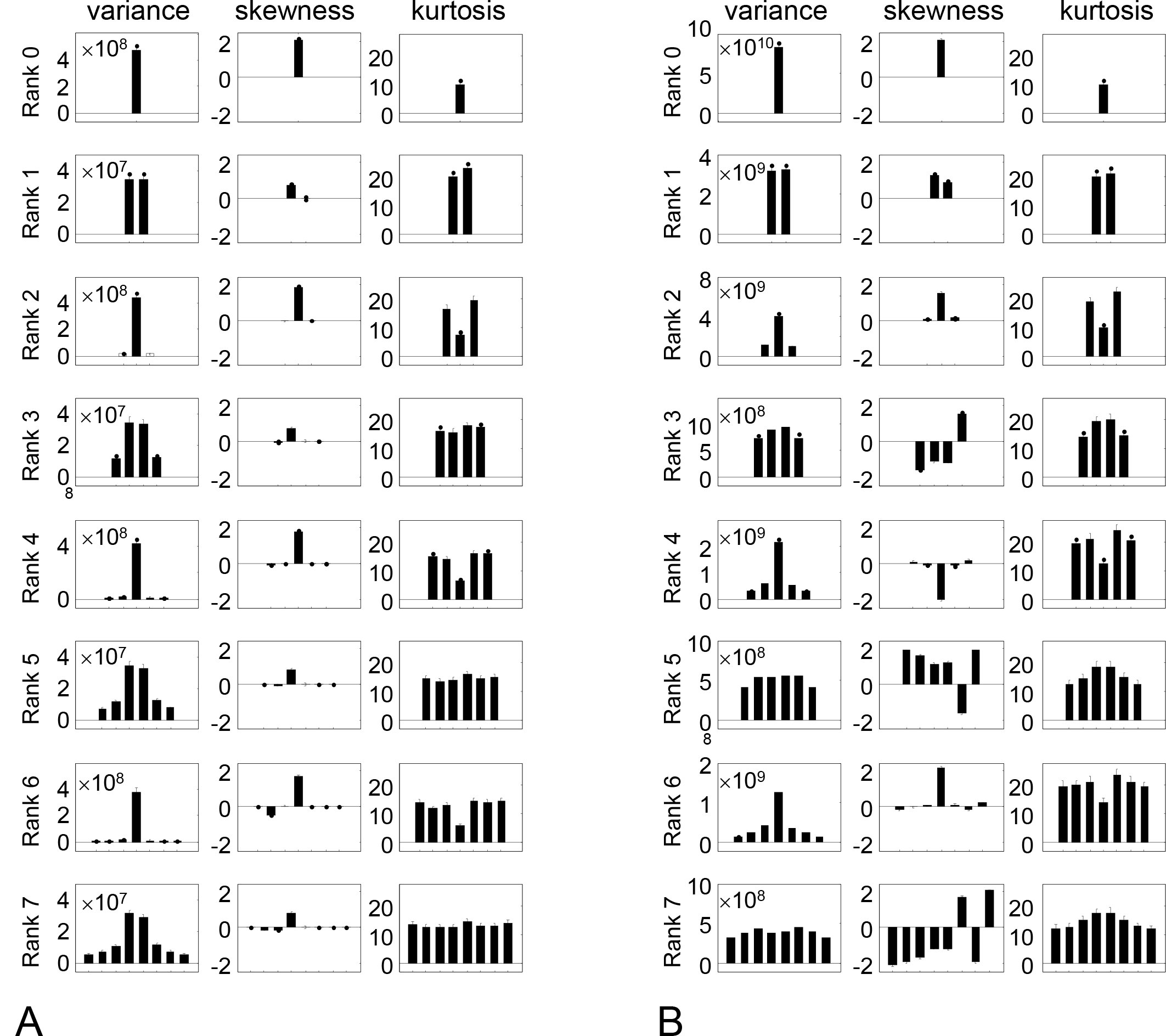
Variance, skewness, and kurtosis for (A) natural images filtered by polar TDH filters of rank 0 to 7 (spatial scale 4), and (B) modified TDH filters in which the polynomial component is replaced by its sign. Error bars are 3 SEM.

With regard to skewness (Figure 6A, second column), there is a single polar filter for which skewness is large; for the others, is close to zero for the others. For even ranks (consistent with the rank-2 results shown in Figure 5), the single polar filter that has a large skewness is the target-like filter 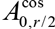; this is the only polar filter with a nonzero mean. For odd ranks, the filter with the largest skewness is the filter with a single horizontal inversion axis, 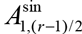; this filter is specifically sensitive to vertical gradients.

With regard to kurtosis Figure 6A, third column, the pattern is also a simple one. For even ranks (also consistent with Figure 2), kurtosis is uniform for all filters except the target-like one 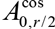, shown as the middle bar of each histogram in the right column); for the target-like filter is approximately half the size of the others. For odd ranks, the kurtosis is large but uniform across all filters. Thus, we find that for each rank, skewness and kurtosis are either uniform across all polar basis functions, or uniform for all basis functions except for one special filter – the odd-rank filter with a single horizontal inversion axis, or the even-rank filter that is target-like.

For completeness, the first column of Figure 6A shows the variance of each filter’s outputs. This is large for target-like filters (center filter in even ranks), and small for all other filters, with sine and cosine pairs resulting in similar variances. As variance is a second-order statistic, this behavior is a consequence of the *k*^-2^ power spectrum of the images.

The simple behavior of skewness and kurtosis for the TDH functions is not merely a consequence of their polar symmetry. To see this, we repeated the analysis of Figure 6A, but with the polar TDH functions replaced by binarized variants, in which positive values of the polynomial component (eq. (3)) are replaced by +1, and negative values by −1. The binarized variants have the same polar symmetry and sine/cosine pairing as the original TDH functions, and, within ranks, are mutually orthogonal as well. However, neither skewness nor kurtosis have the same simple behavior seen in Figure 6A.

While Figure 6A suggests that skewness and kurtosis of a general TDH filter depends only on its projection onto the special axis, it only examines filters that are orthogonal to the special axis. For oblique directions, it is possible that this result will not hold. The reason that more complex behavior may arise in oblique directions is that for moments of order 3 and higher, the steering equations (eqs. (9) and (10) in §2.4) include contributions from mixed moments of the polar TDH’s.

Figure 7 shows that despite this potential complication, skewness and kurtosis of a TDH filter’s output depends chiefly on the projection of the filter onto the single special axis identified in Figure 6A. It is noteworthy that this holds not only for the Cartesian TDH’s, but also for generic TDH’s – which typically lack rotational symmetry. Moreover, for ranks *r* ≥ 3, TDH functions that share the same projection onto this axis are intrinsically different in shape, and are not merely physical rotations of one another.

In sum, within each rank, skewness and kurtosis of the filter coefficient distribution is either uniform, or uniform in all but one direction in filter space. This axis has a simple interpretation – it is either the target-like TDH function, or the single TDH function that is sensitive to a top-to-bottom gradient. In other words, although TDH filter space has a high dimensionality (equal to the rank+1), the behavior of skewness and kurtosis is always low-dimensional – either uniform, or rotationally symmetric.

**Figure 7.**
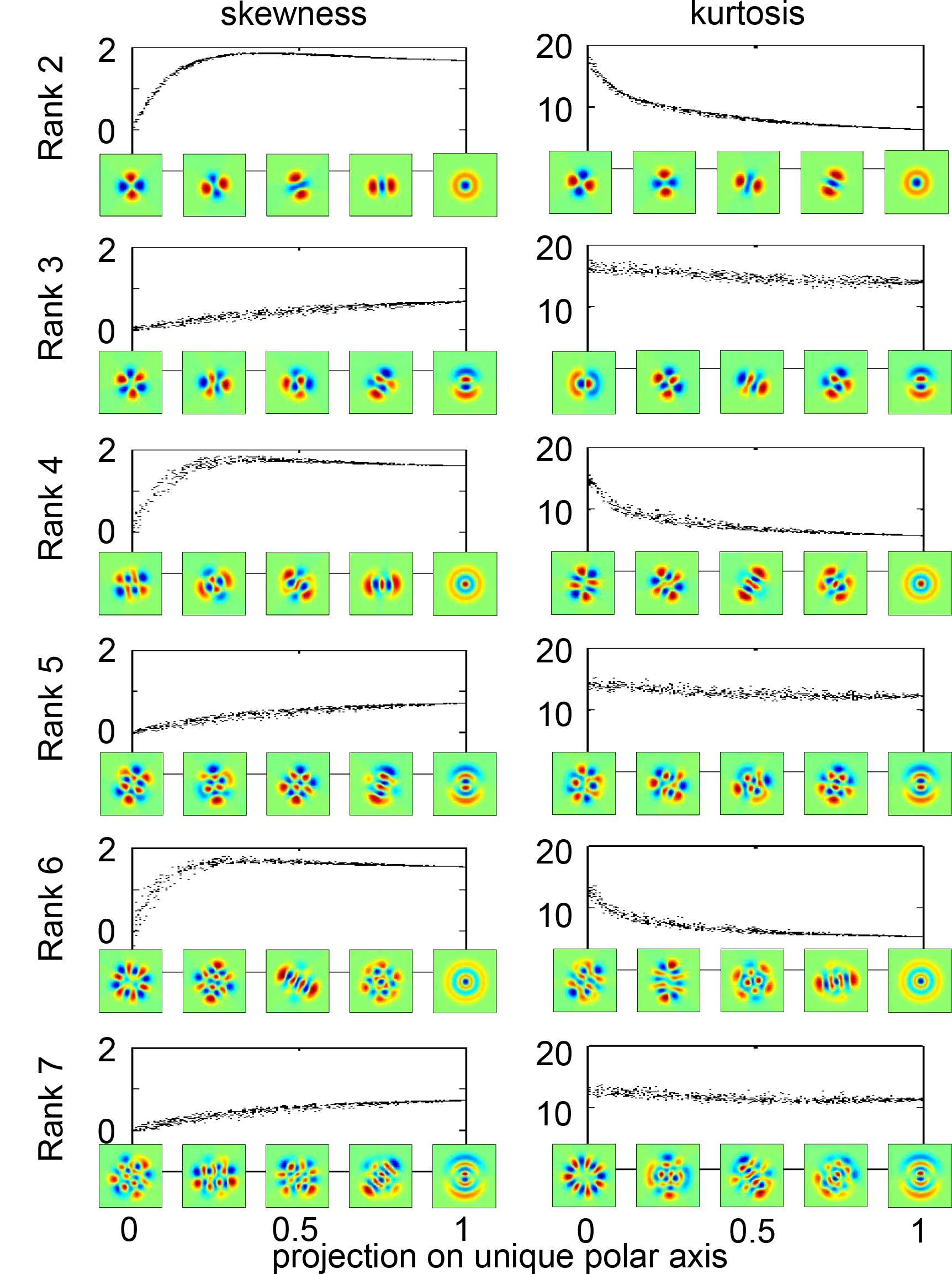
Skewness and kurtosis of natural images filtered by 1000 random TDH filters of rank 2 to 7, at scale 4. The abscissa is the projection of each random TDH filter onto the polar TDH filter shown at the lower right of each plot, which is the target-like filter for even ranks and the filter with a single, horizontal inversion axis for odd ranks. The filters placed along the abscissa are examples of filters whose projections onto the rightmost polar filter are 0, 0.25, 0.5, and 0.75. They illustrate the diversity of filters with a given value of the projection; the examples shown for the skewness and kurtosis columns at corresponding points along the abscissa are interchangeable.

Figure 8 uses this finding to describe the distribution of TDH filter coefficients across all spatial scales in a concise manner. Skewness is characterized by its value for the target-like filter at even ranks (*γ*_3,*target*_, Figure 8A) and for the filter with a single horizontal inversion axis at odd ranks (*γ*_3,*horiz*_, Figure 8B). *γ*_3,*target*_ is a decreasing function of scale and rank and *γ*_3,*horiz*_ is an increasing function of scale, and (except for rank 1) nearly independent of rank. Kurtosis is characterized by its value for the target-like filter at even ranks ( *γ*_4,*target*_, Figure 8C) and by its value for the remaining filters, at both even and odd ranks (*γ*_4,*non-target*_, Figure 8D). Both kurtosis parameters are decreasing functions of scale and rank.

**Figure 8.**
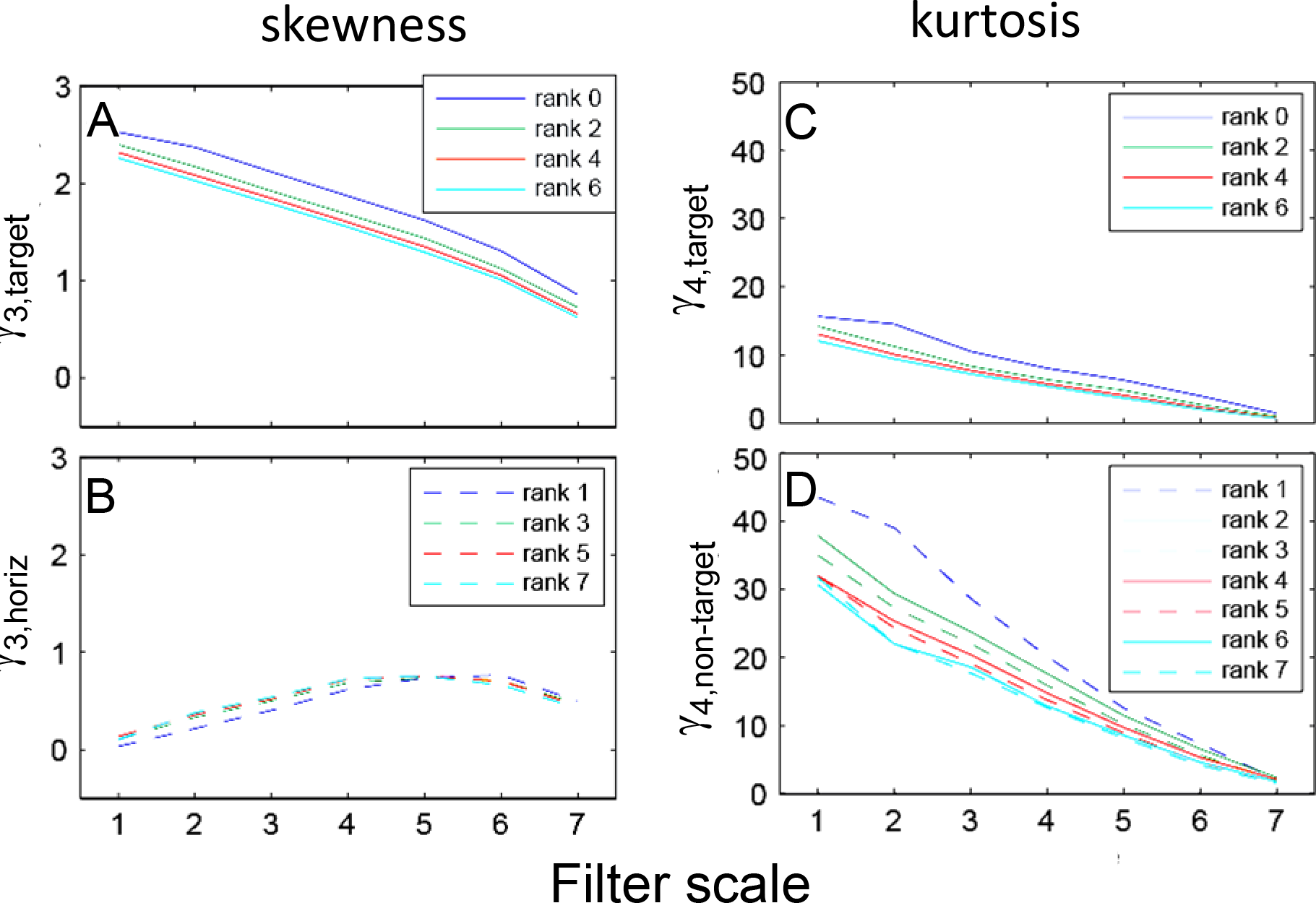
At each spatial scale, skewness of TDH-filtered images is characterized by two values: *γ*_3,*target*_ (A) for even ranks and *γ*_3,*horiz*_ for odd ranks (B), and kurtosis is characterized by *γ*_4,*target*_ (C) for even ranks and *γ*_4,*non-target*_ ranks (D).

### 3.3. Statistics TDH filter coefficients for altered images

To understand the attributes of natural images that underlie the above findings, we carried out parallel analyses for natural images that were manipulated in several ways prior to the determination of filter coefficients.

First, we examined the role of local mean luminance. To do this, we repeated the analysis of Figure 6, but with subtraction of the local mean luminance over a disk of radius 6σ prior to computing TDH filter outputs (Figure 9A). This manipulation eliminated the difference between the kurtosis for the target-like filter and the others, so that kurtosis was uniform within each rank. Subtraction of the local mean reduced, but did not eliminate, the value of the skewness for the target-like filter. As expected, subtraction of the local mean did not change the distributions for the polar TDH filters that were not target-like, since for *μ* … 0, the trigonometric terms in eqs. (1) and (2) necessarily integrate to 0.

**Figure 9.**
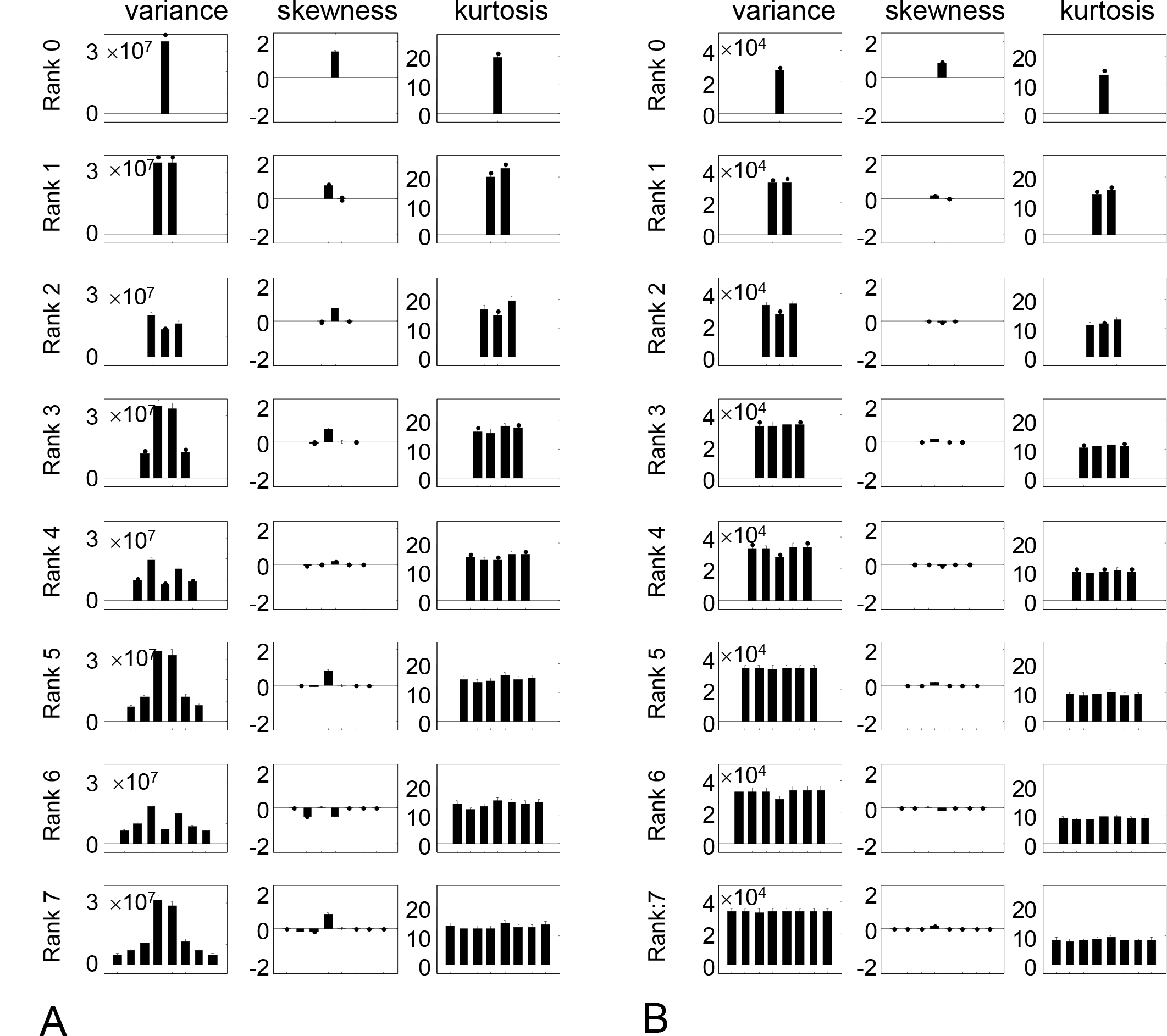
Variance, skewness, and kurtosis for (A) natural images filtered by polar TDH filters of rank 0 to 7 (spatial scale 4) after local mean subtraction. (B) as in (A), but natural images are whitened prior to analysis. Error bars are 3 SEM.

To distinguish the roles of spatial frequency content and phase correlations, we analyzed the distribution of filter coefficients for phase-scrambled images and for images that are spectrally flattened. To isolate the role of spatial frequency content, we created phase-scrambled images by randomizing the phases of the Fourier components in the original images. This effectively results in samples of a spatial Gaussian noise whose power spectrum matches that of the original image. As expected, analysis of these images yielded distributions of TDH filter outputs whose variances matched those of the original images, but for which skewness and kurtosis were zero (not shown). This confirms that spatial frequency content alone does not carry the high-order statistics observed in natural images [8].

To isolate the role of phase correlations, we set the Fourier component amplitudes in the original images to unity, but retained their phases. As in Figure 9A, calculation of filter outputs was carried out with subtraction of the local mean, to retain the isotropy of the kurtosis. Other than for the rank-0 filter, this eliminated the skewness (Figure 9B). The kurtosis remains isotropic. Thus, the heavy-tailed nature of the coefficient distributions depends not only on phase, but also on amplitude.

Finally, to determine the role of the luminance distribution, we calculated the filter coefficient distributions for images subjected to manipulation of the pixel histogram: logarithmic transformation, histogram equalization, and transformation of the intensity histogram to a Gaussian, truncated to 2.56 s.d. (Figure 10). All of these reduced both skewness (by approximately a factor of 10) and kurtosis (by approximately a factor of 5), with near-complete elimination of skewness following the logarithmic transformation. Skewness was concentrated in the filter with a single horizontal inversion axis at odd ranks, and kurtosis was approximately constant within rank.

**Figure 10.**
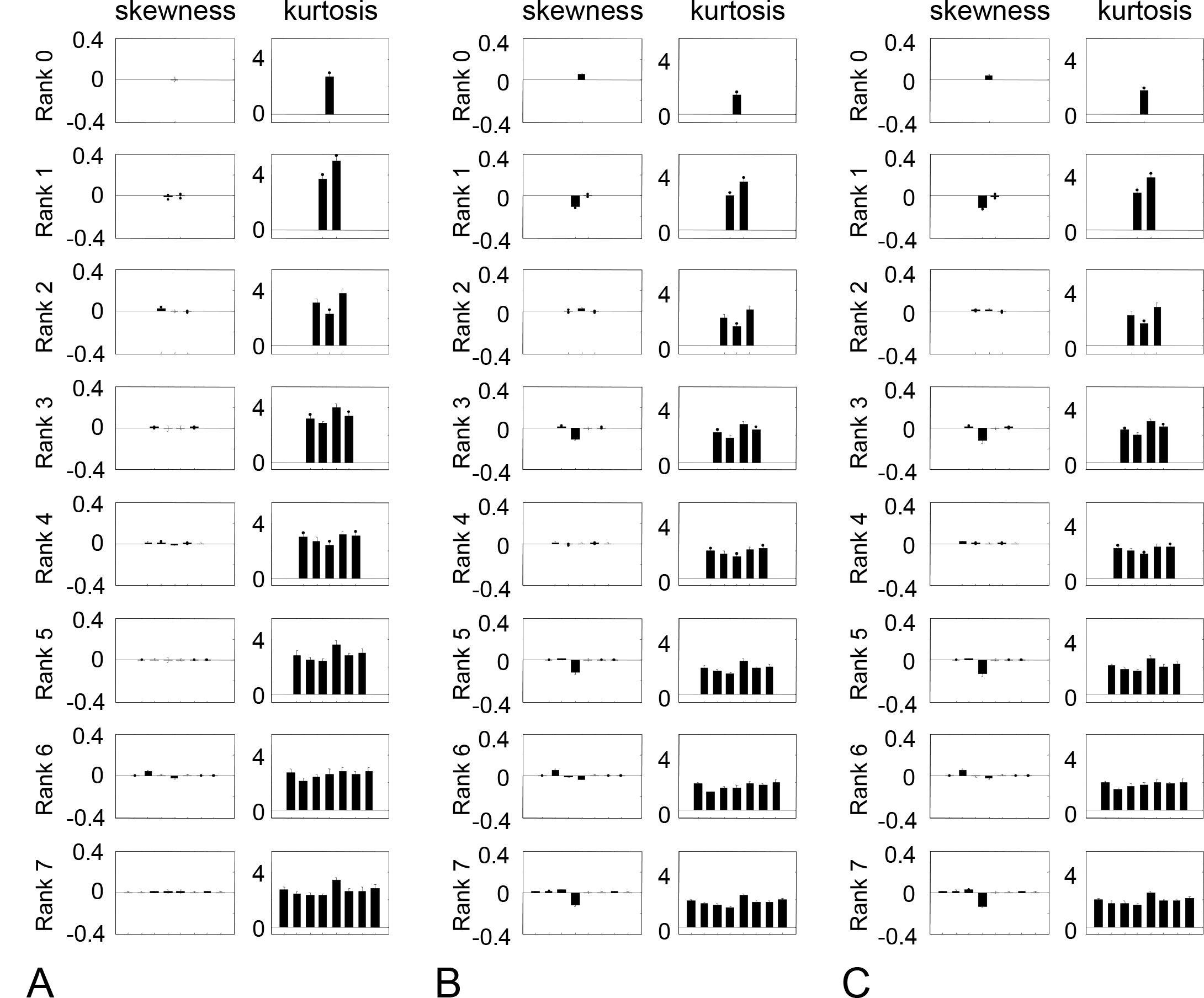
Skewness, and kurtosis TDH filters of rank 9 to 7 (spatial scale 4) processed by pointwise nonlinearities prior to analysis. (A) logarithmic transformation, (B) histogram equalization, (C) Gaussian luminance distribution. Error bars are 3 SEM.

## 4. Discussion

Here we show that two-dimensional Hermite (TDH) filters, an orthogonal basis set with a high degree of symmetry, simplify the description of high-order statistics of natural images, both locally and over wide areas. The significance of this result is that high-order statistics carry the local features that distinguish natural images from Gaussian processes [3, 8, 17, 18, 29], but they are challenging to analyze because of their high dimensionality. By identifying a hidden symmetry in high-order statistics, TDH functions provide a kind of dimensional reduction, and therefore, a needed simplification. We note that this application of TDH functions to characterize natural image statistics is distinct from two other applications of them to vision: a body of work in image processing [19, 21, 23, 24] that uses them to extract local features, and neurophysiologic studies that use them as visual stimuli to analyze the properties of neuronal receptive fields[30, 31].

Our findings can be viewed as building on [17] and [32], which also focus on the high-order image statistics in natural images. Specifically, these authors examined the distributions of outputs of filters acting on whitened natural images, and the joint distributions of outputs of pairs of filters identified by independent components analysis. [17] showed that the joint distribution is approximately circular, and [32] showed that an improved characterization of the joint distribution could be obtained using an *L^p^* -norm, rather than the Euclidean norm. This near-circularity implies that for any filter, the distribution of outputs has a qualitatively similar heavy-tailed shape. However, this is similarity is only a loose approximation: when analyzed quantitatively (e.g., Figure 5 of [17]), the kurtosis of these distributions varied by at least as factor of two. Here, we show that analysis in terms of TDH filters concisely summarizes this variation: at each rank, the kurtosis of a filter’s output is determined by its projection onto a specific direction in filter space.

Examination of the polar TDH filters (Figure 1) suggests the reasons that specific axes are singled out. For the even-rank filters, the special axis is the only filter whose mean is nonzero; all other filters necessarily have a mean of zero because of their sinusoidal dependence on angle. Thus, these filters are the ones that are sensitive to the distribution of local luminances, which are well-known to be heavy-tailed in natural images, both in terms of skewness [33, 34] and kurtosis [35]. For the odd-rank filters, the identified axis has a horizontal mirror-inversion, with large lobes above and below the horizon. Thus, these filters are likely to be highly sensitive to vertical gradients, and thus, the distributions of their outputs will be skewed by the tendency of illumination to come from above. Consistent with these hypotheses, removal of the local mean (Figure 9A) eliminated the distinctive behavior of target-like filter for kurtosis, and reduced its skewness. When the low-spatial frequencies were reduced by spectral flattening, the skewness was eliminated for the odd-rank filters as well. Figure 10 provides further evidence that the distinctive kurtosis for the target-like filters is primarily a consequence of luminance distributions, as it is reduced by attenuating the tails of the luminance distribution via log transformation, histogram-equalization, or Gaussianization.

However, the simplification we observe is not simply a consequence of the arrangement of the positive and negative lobes of the TDH filters. The evidence for this is that replacing the Hermite polynomial values by ±1, which preserves the arrangement of their lobes, does not result in a similar simplification of the skewness and kurtosis (Figure 6B). Thus, the gradations of the polynomials that define the TDH’s, which underlies their generalized steerability, is crucial.

## 5. Conclusions

Two-dimensional Hermite filters provide a simple description of third-and fourth-order statistics of natural images across a range of scales. This simplification is a consequence of the high degree of symmetry of this orthogonal basis set, and the phase, amplitude, and luminance characteristics of natural images.

## Acknowledgments

We thank Eyal Nitzany and Matthias Bethge for comments on an earlier version of this manuscript. Supported in part by NIH EY07977 and NIH EY09314 to J.V.

## Author Contributions

J.V. and Q.H. designed the experiments; Q.H. carried out the analysis; J.V. and Q.H. wrote the paper.

## Abbreviations

The following abbreviations are used in this manuscript:

TDH: Two-dimensional Hermite

